# Speed of time-compressed forward replay flexibly changes in human episodic memory

**DOI:** 10.1101/323774

**Authors:** Sebastian Michelmann, Bernhard P. Staresina, Howard Bowman, Simon Hanslmayr

## Abstract

Remembering information from continuous past episodes is a complex task. On the one hand, we must be able to recall events in a highly accurate way that often includes exact timing; on the other hand, we can ignore irrelevant details and skip to events of interest. We here track continuous episodes, consisting of different sub-events, as they are recalled from memory. In behavioral and MEG data, we show that memory replay is temporally compressed and proceeds in a forward direction. Neural replay is characterized by the reinstatement of temporal patterns from encoding. These fragments of activity reappear on a compressed timescale. Herein, the replay of sub-events takes longer than the transition from one sub-event to another. This identifies episodic memory replay as a dynamic process in which participants replay fragments of fine-grained temporal patterns and are able to skip flexibly across sub-events.

## Introduction

Episodic memory retrieval is a flexible process that operates at different timescales ^1^. In some instances, it is crucial for our behavior to mentally replay events at the same speed as the initial experience: Re-enacting a classic movie scene relies on a temporally accurate representation of dialogue and events. In other instances, it would be highly dysfunctional to recall our memories at the same speed they originally unfolded: We have to be able to reconstruct how we came to work today without zoning out at our desk for thirty minutes and must therefore be able to flexibly adjust the speed of our memory replay.

Previous work has already put forward that memory replay in humans could be forward and compressed: Studies that related the timescale between retrieval and perception of a particular event asked participants to mentally navigate routes based on their memories. The duration of memory replay (i.e. mental navigation) was found to be faster than the real navigation, but varied substantially between participants ^2,3^. Interestingly, findings of neural replay in rodents mirror this compression, by showing that hippocampal place cells, which correspond to certain positions along the animal’s path, later fire again on a faster timescale than during navigation. This is interpreted as reflecting compressed replay of past trajectories ^4,5^. One recent study in humans observed the reactivation of static representations in electrocorticography (ECoG) and found patterns of oscillatory gamma power reappearing faster than during perception ^6^. On the other hand, several studies find reappearing temporal patterns from perception in memory, demonstrating that some patterns are replayed at the same speed ^7–10^. Notably, a recent study that applied functional magnetic resonance imaging (fMRI), managed to track continuous memory reinstatement over long episodes (50min). Spatial patterns reappeared during the free recall of narratives; recall was temporally compressed, but varied between participants ^11^.

Despite these indications about the speed of replay in behavioral and neural data, no study so far has tried to directly read out the temporal dynamics of memory replay in humans on a fine-grained temporal scale. Importantly, the recent advent of multivariate methods in neuroscience has now opened new avenues for the investigation of these processes: By leveraging multivariate patterns in combination with electrophysiology, it is now possible to track representations from perception in a time-resolved manner, as they reappear during memory retrieval ^6–8,12–15^. Importantly, simultaneous EEG and multi-unit recordings in primates demonstrate an intimate relation between neural firing and the phase of slow oscillations in the EEG ^16^. Therefore, information about neural patterns can be captured and tracked in human electrophysiology via oscillatory phase ^8,16,17^.

Capitalizing on these methodological advances, we here investigate the flexible dynamics of episodic memory replay in continuous mnemonic representations. Studying these trajectories during memory replay requires a paradigm that prompts participants to evoke continuous representations with distinct subevents from memory. This will make it possible to track fragments of these representations in episodic memory via multivariate analysis methods. To this end, we asked subjects to associate static word-cues with ‘video-episodes’ consisting of a sequence of three distinct scenes. The three dynamic scenes thus formed a continuous six-second-long video. In encoding-trials, we presented a word-cue during one of the scenes. This allowed us to prompt memory replay in a natural way, i.e. we asked participants to recall in which of the three scene-positions they had learned an association during encoding. After completing this part of the task, we asked about the video-episode itself and confirmed memory accuracy. In a behavioral experiment, we investigated direction and speed of replay via measuring reaction times to the scene-position response. In a separate MEG study, we leveraged the content-specific phase patterns that each scene elicited and used them as handles to track the direction and speed of replay of the video-episodes. If memory replay were indeed compressed, we expected to find evidence for this compression in reaction times and in the reinstatement of neural patterns. This replay could either be forward or backward. In line with previous findings, we expected to find evidence for reactivation of temporal patterns, signifying replay at the same speed for fragments of neural activity ^7–10^. We further hypothesized that the disparity between accurate representations and overall compression would be due to a flexible mechanism that allows subjects to skip between sub-events, as they replay episodes. Therefore, replay within sub-events should occur at a slower pace, whereas skipping between sub-events should occur fast.

## Results

### Compressed and forward memory-replay in reaction times

In the behavioral experiment, participants associated word-cues with one of three scenes within video-episodes (Fig. 1a). We used four continuous video-episodes, each consisting of three individual dynamic scenes. A trial-unique word-cue appeared in one scene during a video-episode. After a brief distractor task (Fig. 1b) subjects performed, in alternation, either a cued-recall (CR) retrieval task or an associative-recognition (AR) task (Fig. 1d, top). The AR task was included as a control condition, because active replay is arguably not required for recognition. In the CR blocks, we presented participants with the word-cues (Fig. 1d, top-left). Their task was to recall the scene-position that was associated with the word-cue as quickly as possible. In AR blocks, subjects successively saw the word-cues superimposed on screenshots from encoding and were asked to decide as quickly as possible whether this association was intact or rearranged (Fig. 1d, top-right).

To address the direction and speed of memory replay, reaction times (RTs) at retrieval were compared between associations that were learned in the first, second and third scene-position of a video-episode (Fig. 1d, bottom). We only used RTs for correct hit trials (correct recall in CR and correctly recognized intact associations in AR blocks) and excluded trials in which the subjects were wrong or guessed (see Supplemental Information for the same analysis including correct guesses). If the CR condition indeed elicited replay in the forward direction, we should observe faster reaction times with CR, but not AR, for associations that were learned earlier during encoding. Furthermore, if replay was compressed, the delay between reaction times to different scene-positions should be smaller than the duration of the scenes segments themselves. The resulting 3x2 repeated measures ANOVA tested the factors position and condition. A significant main effect of position (*F*_1.85, 42.48_ = 5.884, *p* = 0.007, log-RT: *F*_1.79, 41.26_ = 3.375, *p* = 0.049) and a position by condition interaction (*F*_1.75, 40.34_5.9, *p* = 0.008, log-RT: *F*_1.76, 40.58_ = 5.606, *p* = 0.009) were obtained. Both effects were driven by the cued-recall condition (ANOVA position only: *F*_1.79, 41.19_ = 9.082, *p* = 0.001, log-RT: *F*_1.60, 36.90_ = 8.207, *p* = 0.002): During encoding, individual scenes of each video-episode lasted 2 seconds. During CR retrieval, however, associations that were learned in the first scene-position of a video-episode (mean RT = 2.5s) were recalled on average 116ms faster than associations that were learned in the second scene-position (*t*_23_ = −1.870, *p* = 0.037, log-RT: *t*_23_ = −2.4, *p* = 0.012). Associations that were learned in the second scene-position (mean RT = 2.617s) were recalled on average 176ms faster than associations that were learned in the third scene-position (*t*_23_ = −2.767, *p* = 0.006, log-RT: *t*_23_ = −2.274, *p* = 0.016, (mean RT = 2.793s)). The replay of the video-episodes was therefore forward and compressed during CR, which replicated our findings from a behavioral pilot experiment (see Supplemental Information). The average RT difference of 146ms per position corresponds to a compression factor of 13.7 during replay.

Might the effects be due to asymmetrical encoding of scene-positions? One could argue that associations have a higher saliency when they are presented in the first scene-position, leading to higher confidence and shorter RTs during retrieval. Additionally, subjects can take more time to rehearse early associations during the remainder of the video-episode, perhaps resulting in the weakest memory trace for the last scene. Importantly, however, if the serial position merely affects the overall strength of the memory trace in our paradigm, we should observe comparable effects on cued recall (CR) and associative recognition (AR). Conversely, if the effect is contingent on the need to mentally replay scene after scene, serial position at encoding should only exert an effect on the CR task.

Importantly, no differences in reaction times between scene-positions were evident in the AR task (Fig. 1d, right; ANOVA: *F*_1.64, 37.66_ = 0.708, *p* = 0.472, log-RT: *F*_1.61, 36.95_ = 0.793, *p* = 0.435, pairwise comparisons of positions: all *p*s > 0.199, all *BF*_01_ > 2.158). Together with the significant position by condition interaction, this confirms that the position effect on RTs is specific to the CR task and rules out a saliency-based explanation. Finally, we observed a significant main effect of condition with unscaled (*F*_1.00, 23.00_ = 62.349, *p* < 0.001) and log-transformed (*F*_1.00, 23.00_ = 95.036, *p* < 0.001) reaction times. This was due to faster RTs in associative-recognition blocks (*t*_23_ = −7.896, *p* < 0.001, log-RT: *t*_23_ = − 9.7487, *p* < 0.001). Taken together these results are evidence that successful recall of elements from a continuous video-episode relies on compressed forward replay.

**Fig. 1.**
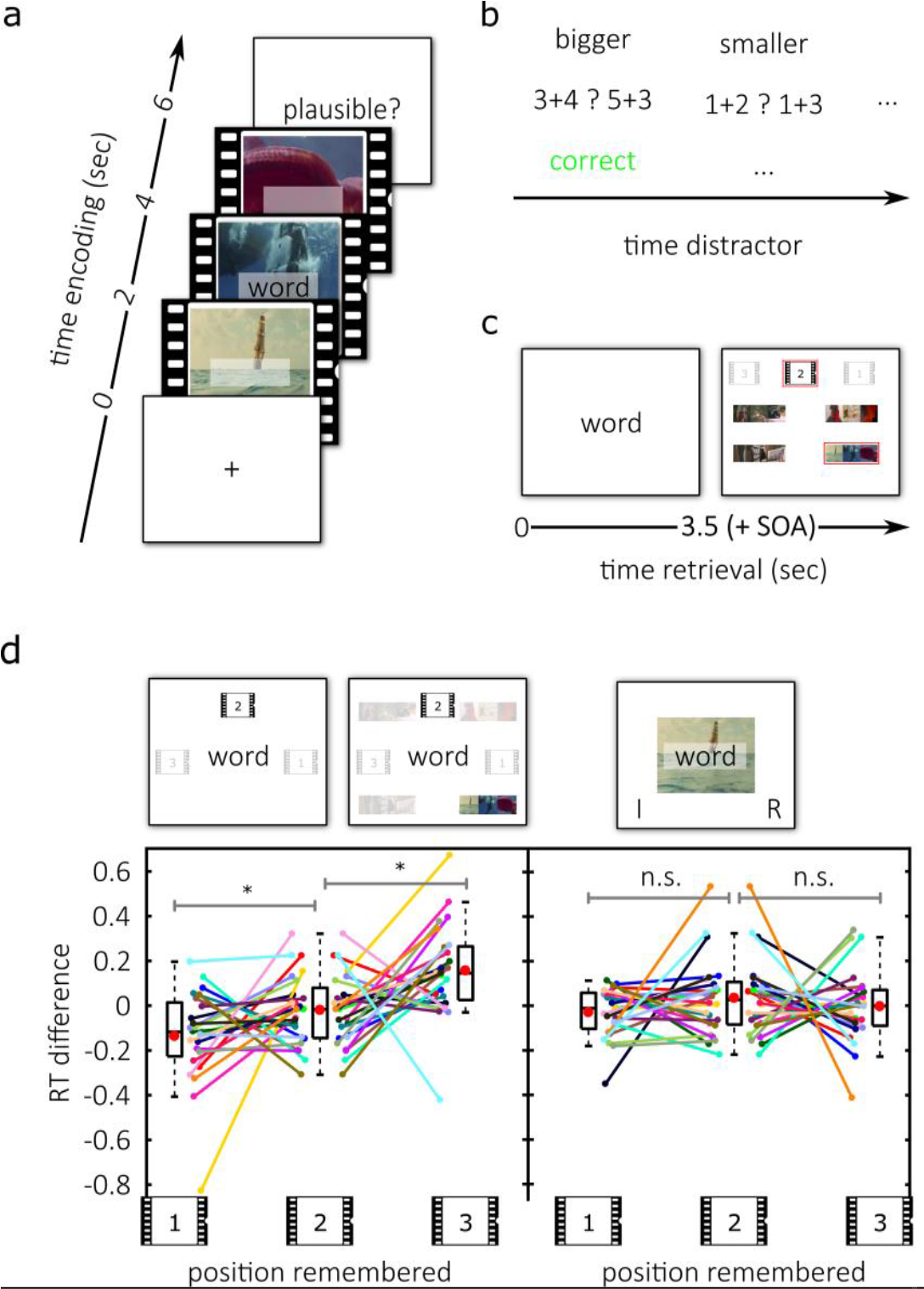
Experimental design and behavioral results. **(a)** During encoding subjects repeatedly saw one out of four video-episodes. In one of three scenes that comprise a video-episode, a word-cue appeared in the center of the screen. (**b)** In the distractor block participants identified either the bigger or the smaller one of 2 simple sums. (**c)** In the MEG experiment, participants saw the static word-cue during retrieval for 3.5 seconds, followed by a fixation cross for 250ms - 750ms. Subsequently they first picked the scene-position in which they learned the association and then confirmed the correct video-episode. **(d)** In the cued-recall (CR) condition of the behavioral experiment (left) participants selected the correct scene position as quickly as possible during retrieval. In an associative-recognition (AR) control condition (right) they decided whether the presented association (word superimposed on a screenshot) was intact or rearranged. In CR blocks, subjects were faster to recall an association that was learned in earlier scene-positions during encoding (bottom left). Importantly, in the control condition, they performed the same encoding task and needed source memory for AR retrieval, however no modulation of reaction times was found. The y-axis denotes the difference to each participant’s average reaction time in the respective condition. Spaghetti-plots show individual subjects. Boxplots are 25^th^ and 75^th^ percentile and the median; whiskers are maxima and minima, excluding outliers. Red dots within the boxplots depict the arithmetic mean. Significant differences are marked with a star, n.s. denotes non-significant in a post-hoc paired t-test comparison.

### Low frequency phase patterns from encoding reappear during successful memory retrieval

In the MEG experiment, participants performed the same CR task as in the behavioral experiment, with the only difference being that they gave responses after the word-cue disappeared (Fig. 1c). In a first step, we asked whether perceptual content could be distinguished based on oscillatory phase. To this end, we compared the inter-trial phase coherence (ITPC) between encoding-trials grouped according to their video-content against the ITPC between trials grouped randomly. This has been used previously to reveal the content specific entrainment of cortical rhythms to naturalistic dynamic stimuli ^8,16^. The four video-episodes showed reliably distinguishable phase patterns during encoding (*p*_cluster_ < 0.001, Fig. 2a, left and middle). The significant cluster (across time space and sensors) contained robust differences in the lower frequencies and showed a maximum over occipito-parietal sensors (Fig. 2a, middle). Consistent with our previous results ^8^, strongest differences were observed at the onset of each scene. Importantly, the frequency band centered at 8 Hz was included in the cluster, which was previously linked to the reinstatement of phase patterns ^8^. Testing the 8 Hz phase differences on the source level revealed one broad cluster of content specificity during encoding (*p*_cluster_ < 0.001). Averaging t-values across this significant cluster over time revealed highest values in occipital and parietal locations (Fig. 2a right). Together, these results show that every sub-scene within a video-episode was associated with a content specific fingerprint in oscillatory phase, which was maximal in a parieto-occipital region. In the following, we used these sub-scene specific phase patterns at the center frequency of 8 Hz as handles to track replay in memory.

In a first step, we tested whether these phase-patterns of the video-episodes were reactivated in memory. Therefore, we first contrasted phase-similarity between encoding-retrieval combinations of the same video-episodes (e.g. watching video A, recalling video A) with encoding-retrieval combinations of different video-episodes (e.g. watching video A, recalling video B). Similarity between encoding and retrieval phase patterns was analyzed with a sliding-window approach (window size = 1 sec), providing a time resolved measure of memory replay ^8,18,19^ (see Fig. 2c). On the source level, analysis was restricted to an anatomically defined occipito-parietal region of interest (ROI) following the results from the encoding phase and previous studies showing memory replay in these regions^8,20–23^ (Fig 2b). Evidence for replay was found for hit trials (Hits; *p_cluster_* = 0.034; Supplemental Fig 3a, also see Supplemental Fig 3b for unmasked maps of t-values), suggesting that replay of video-episodes can be tracked in the phase of an 8Hz oscillation. Notably, we found no such replay effect for Misses, i.e. trials in which subjects either guessed, or did not remember the correct scene-position and/or video-episode. Furthermore, a direct contrast between Hits and Misses revealed significantly stronger replay for Hits compared to Misses (*p*_*cluster*_ = 0.030, Fig. 2d), demonstrating the functional significance of this pattern-reinstatement for memory.

**Fig. 2:**
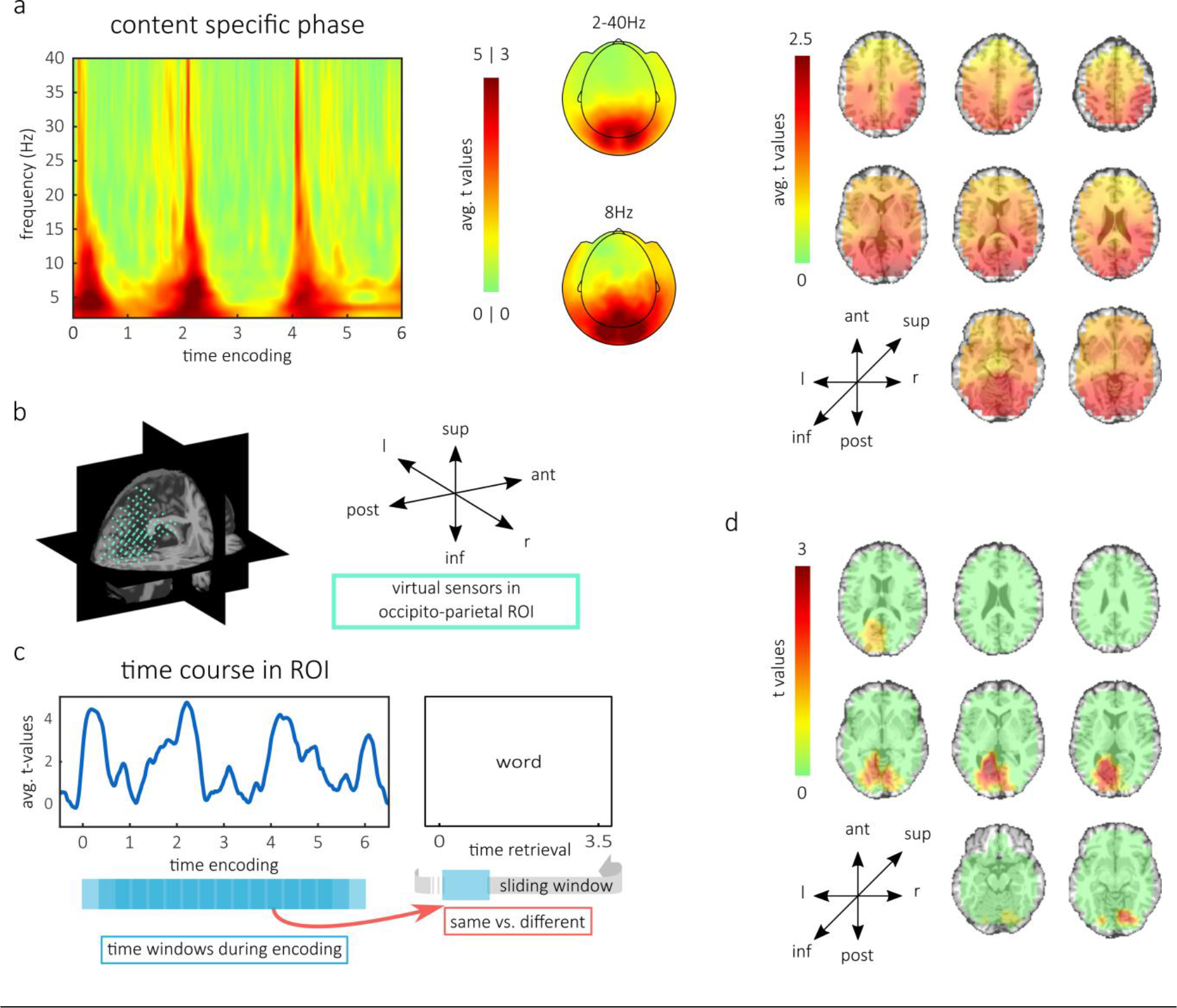
Reinstatement of oscillatory patterns from encoding. **(a)** During encoding, the different video-episodes elicited content specific phase patterns. The left panel shows the averaged t-values across sensors in the cluster of significant content-specificity. Topographies in the middle are t-values within the same cluster (across time, frequency and space), averaged across time and across all frequencies (top) or only for 8 Hz (bottom). Both topographies show maximal values over occipital and parietal sensors. The right panel shows the average t-values across time on virtual sensors, within the temporo-spatial cluster of significant differences at 8Hz. Occipital and parietal sensors expressed the maximal t-values. **(b)** Occipito-parietal region of interest (ROI) that we used for statistical testing of content-specific reactivation. **(c)** Time course of content specific phase during encoding, averaged across the ROI. Below, the sliding window approach is illustrated, in which all possible time windows from encoding were compared to each retrieval time window via phase coherence. Subsequently, combinations of same and different content combinations were contrasted. **(d)** Cluster of significant differences between content-specific reactivation for successfully remembered and forgotten associations.

### Compressed replay is forward

The above findings confirm that content specific patterns of activity from encoding, are reinstated in a purely memory driven way. This motivated us to ask in which direction and at what relative speed patterns from encoding unfold during retrieval. Do patterns from the beginning of the video-episodes, for instance, also reappear earlier during memory retrieval?

To this end, we divided the encoding interval into 6 non-overlapping windows, centered at 0.5, 1.5, 2.5, 3.5, 4.5, and 5.5 seconds. We then analyzed the phase-similarity to these windows across the retrieval interval. The latency at which patterns reappear should be reflected in the distribution of phase-similarity across time. Consequently, we compared these distributions between the distinct time windows from encoding (Fig. 3a).

Specifically, to test the direction of replay statistically across subjects, we used the following approach: We cumulated the similarity distributions across the whole retrieval time. This provided the cumulated similarity (CS) for every subject and every encoding-window. Similarity started at the beginning of the retrieval interval with a value of zero. It ended at the end of the retrieval interval, with a value of one (Fig 3c). If phase-similarity to an encoding-window “A” cumulates earlier than phase-similarity to an encoding-window “B”, then the cumulated similarity for “A” is higher compared to “B” and consequently “A” is replayed earlier during retrieval than “B”. In other words, when the CS of one phase-pattern is higher than the CS of another, then the evidence for replay of that phase-pattern is leading over the other at that point. If, however replay of a phase-pattern is lagging behind the replay of another, the CS should be lower at that time point. We tested this relation statistically at every time point by comparing the cumulated similarity across all windows for each subject. The overall tendency is tested best by fitting a line across all six encoding windows. Herein, a negative slope indicates forward replay, since earlier windows have higher values in the CS than later windows, a positive slope signifies backward replay.

Results revealed significant forward replay in two time windows (i.e. 135ms to 1919ms, and 3458ms to 3473ms after cue presentation, see Online Methods for some notes of caution regarding the interpretation of the exact time-window). We can therefore conclude that there is a dominance of early encoding-patterns in early time points at retrieval, relative to late encoding-patterns. This supports the notion of forward replay (see also Supplementary Information for additional evidence supporting forward replay) and corroborates the finding of forward replay from the behavioral experiment.

**Fig. 3:**
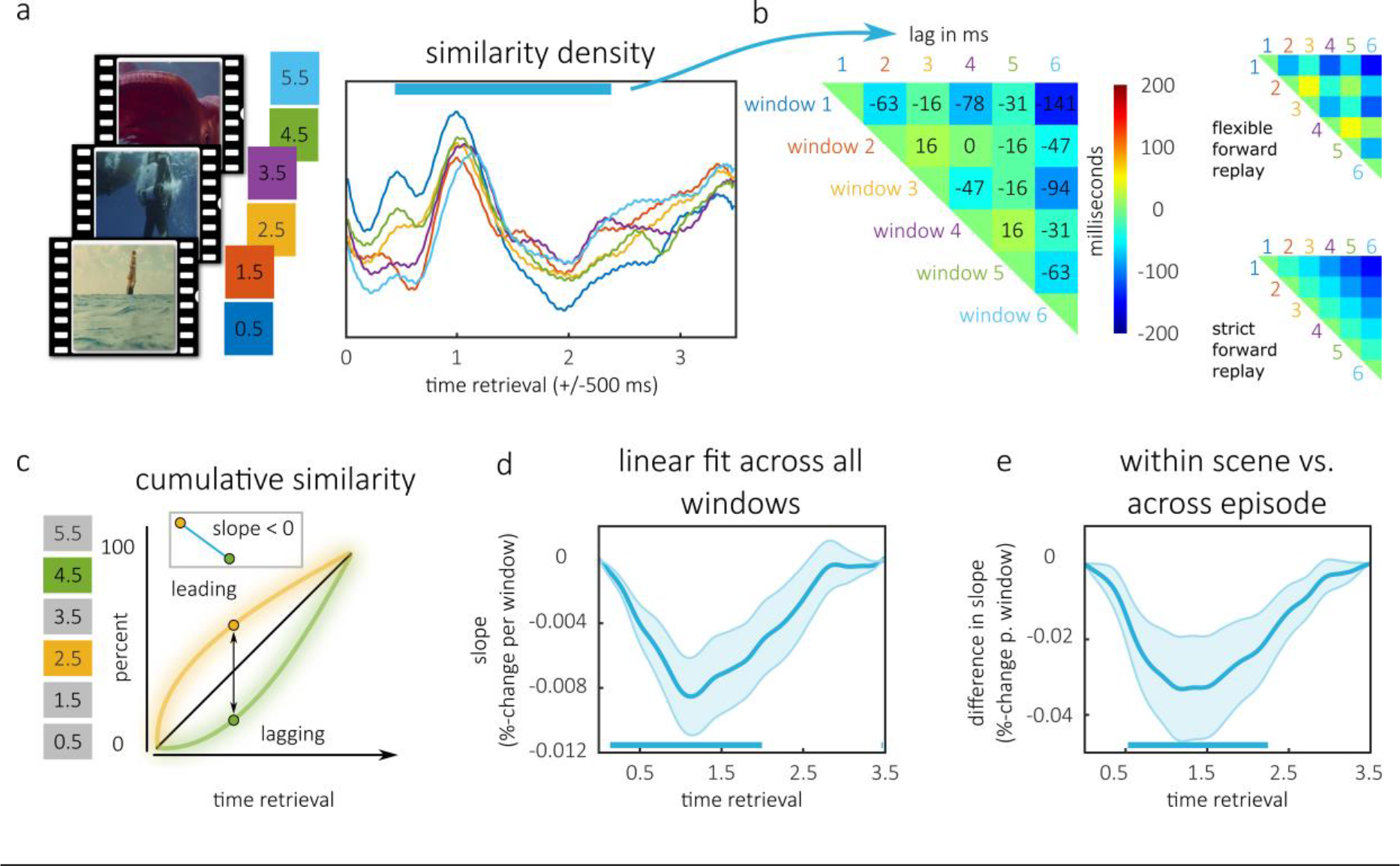
Chronometry of memory replay. **(a)** The 6 non-overlapping time windows from encoding illustrated next to a video-episode (left). The average similarity densities to these windows are on the right. The blue bar denotes where replay was significantly slower within scenes (see e). **(b)** Cross correlations of similarity densities within this window show the adaptive pattern. The matrix shows the combination of windows that are correlated in each cell. The times in ms at which cross correlation is biggest are displayed in the color-coded cells. In this, lags between windows within scenes are bigger than lags between windows across scenes (right, top); with strict forward replay, all scenes would be replayed in order (right, bottom). **(c)** Illustration of the cumulative similarity (CS) approach used to test replay-dynamics. If evidence for a window statistically precedes evidence for another during retrieval, its cumulated similarity is higher. Fitting a line through those subsequent encoding windows will therefore result in a negative slope. **(d)** Average slope of lines fit across all windows’ CS, for each subject and time point. Negative slope indicates that earlier encoding-windows have higher CS values and signify forward replay. **(e)** Contrast of average slopes from the average fit across windows within scenes and a fit across all windows, supporting an adaptive replay framework. The blue bars in d and e denote significance.

### Speed of replay is flexible

In neural data, the forward direction of replay was evidenced by the tracking of content specific temporal patterns. Notably, however the reactivation of temporal patterns signifies that participants replayed *fragments* of the video-episodes at roughly the same speed as during encoding. Hence, these data already indicate that memory replay is not the straightforward recapitulation of the original experience. Instead, flexible processes must be at work to reconcile the overall compression of memory with the reappearance of temporal patterns.

We hypothesized that the disparity between locally detailed patterns and the global compression was possible through the flexible skipping between salient components in memory (e.g. sub-events); in our data, the boundaries between scenes were salient elements within the video-episodes. We therefore investigated whether these boundaries would serve as stepping stones enabling participants to skip through their memories on a faster time-scale. Consequently, we tested statistically whether the speed of replay slowed down within scenes, since the skipping between the scenes of a video-episode should be easier and more likely than skipping within the individual scenes.

To this end, we extended the method of fitting a line across CSs to compare the compression of replay within individual scenes (i.e. within sub-events) to the overall compression level. Specifically, calculating the slope of the fitted line allows for an estimation of the speed of replay. This slope indicates the lag between replayed patterns in the retrieval interval, such that steep slopes indicate a long lag (i.e. slow replay). We fitted a separate line for each pair of encoding-windows that belonged to the same scene across their respective CSs and averaged the slopes across the three lines. The time interval between 442ms and 2350ms displayed slopes significantly below zero, confirming forward replay within scenes. More importantly, between 550ms and 2350ms at retrieval, slopes of windows within a scene were significantly steeper (i.e. replay was slower) compared to the slope obtained across all encoding-windows (Fig. 3e). This means that when participants replayed the first and second part of a scene, this replay was less compressed than we expected from the global compression level of the whole video-episode. Consequently, this also means that subjects did not recapitulate every scene successively in every trial. Taken together, these results show that memory replay does not occur at a constant speed; instead, the speed of replay seems to change flexibly depending on the replayed interval (Fig. 3b, right). Finally, we repeated these tests with those trials in which subjects did not remember the correct positional-scene or video-episode; however, we found no significant time-points for any of the contrasts, which demonstrates the implication of these replay effects in memory. In a further control analysis, we excluded the first 800ms of the retrieval interval for the similarity analysis in order to rule out that event related potentials (ERPs) drove similarities. Again, we found significant negative slopes between 812ms and 1212ms and slower replay within scenes in that window.

These results statistically support a flexible forward replay strategy. Via cross-correlations, we next derived a descriptive measure of the delay between the six sub-events during flexible memory replay (550ms–2350ms). The cross-correlation was computed on pairs of averaged and smoothed similarity distributions (Fig. 3b), which retained a time lag value for every combination of the six sub-events. The adaptive replay that we found is also visible in the pattern of time lags and can be illustrated with shorter lags between time windows that belong to different scenes compared to time windows that belong to the same scene (Figure 3B, right). In contrast, to illustrate a strict and inflexible forward replay strategy, lags between the sub-events should increase linearly according to their position at encoding (illustrated in Figure 3B, right).

## Discussion

In this study, we tracked the replay of continuous episodes from memory. We used a novel paradigm in which participants associated unique word-cues with one out of three distinct scenes in seamless video-episodes. We prompted replay by asking volunteers in which exact position (1, 2, or 3) they had learned each word-cue. Behavioral and neural data indicated that replay of memories takes place in a forward direction and at a compressed speed, i.e. memory replay was faster relative to perception. Notably, on a neural level, we found indications for different speeds of replay: Fragments of temporal patterns reappeared at the same speed and the speed of replay within sub-events (i.e. scenes) of continuous video-episodes was slower than the overall compression level.

Importantly, our finding of different compression levels implies that memory replay acts in a flexible way. The disparity between the slower speed of replay within scenes and the overall compression is an aggregated observation that cannot hold on a single trial level. Specifically, it signifies that replay is not a simple concatenation of fragments because in a single trial, the sequential replay of three scenes would take longer than the overall compression permits. Consequently, participants must be able to skip between replayed fragments; importantly the slower speed of replay within scenes denotes that on average, the skipping between sub-events must take place on a faster temporal scale than the skipping within sub-events. A plausible interpretation of the observed pattern is therefore that replay of relevant information is initiated from the boundaries between scenes and that participants can flexibly skip between them. Event boundaries ^24^ have been previously shown to trigger replay events during memory encoding ^25^. They could therefore also serve as starting points during memory retrieval, to initiate the replay of information on a fine-grained temporal scale. Mechanistically, the hippocampus has been suggested to preserve the temporal order of experiences ^26^ and interactions between the hippocampus and visual cortex have been observed during memory replay in sleeping rodents ^20^. In our data, we consistently found reinstatement of fine-grained temporal patterns in sensory-specific regions ^8^. It is therefore possible that the hippocampus exerts control over sensory areas, when those regions execute the vivid reinstatement of sensory information. Specifically, information-rich and temporally accurate representations could rely on sensory cortices whereas the hippocampus initiates replay, based on a sparse code ^27^. At first glance, the reinstatement of temporal patterns is also at odds with the observation of compression in general. An important implication from the finding of temporal pattern reinstatement under global compression is therefore that the accurate reinstatement of patterns must be limited to fragments of the original perception. In other words, subjects possibly omit non-informative (perhaps redundant) parts of the video-episodes and therefore replay a shorter episode in memory, which contains less information. Previous work on mental simulation of paths supports this interpretation. The duration that participants take to mentally simulate a path increases, when this path includes more turns ^3^. In the same way, the duration of replay might depend on the overall number of relevant elements within a video-episode.

Another crucial result from our experiments is the forward direction of replay. This finding is in line with recent studies showing anticipatory activation of familiar paths in the visual cortex ^22^ and evidence of forward replay of long narratives ^11^. Notably, in the rodent literature, the task of spatial navigation appears to determine whether replay is backward or forward. At the end of a path, awake rodents replay in a backward fashion ^4^, whereas animals that plan the path towards a goal display an anticipatory activation of place-cells in the forward direction ^28^. Task requirements in our design could indeed have prompted participants to step mentally through the video-episodes in a forward manner. Speculatively, other designs (e.g. tasks requiring recency judgments) might therefore cause a backwards replay. This would be well in line with the flexibility in memory replay that we observed in the neural data, since a flexible mechanism could arguably guide replay in a forward and backward direction when skipping through events. An interesting additional question arising from this is whether replay of fine-grained temporal patterns in the cortex can also be backwards.

Importantly our study also demonstrates how one can investigate these open questions. The design that we used to trigger the replay of distinct sub-events in a continuous episode can easily be adapted to a working memory context and our method to track oscillatory patterns allows for the investigation of replay in working memory, during rest and during sleep. We have repeatedly shown how to use the similarity in oscillatory phase to track content-specific reactivation, even when the exact onset of memory-reactivation is unknown. We here extended our previously developed method ^8^ to track distinct sub-events from continuous representations: In a statistically robust way we aggregated evidence across several repetitions and compared their distribution across time.

This investigation of temporal dynamics during human episodic memory replay has only recently become an option, when the tracking of multivariate patterns was extended to human electrophysiology ^7,8,10,14,15,25,29^. Leveraging a novel paradigm in combination with a method that can detect the individual fingerprints in oscillatory patterns, we were now able to observe the fine-grained dynamics of memory replay on a behavioral and on a neural basis. Our data render memory replay as a flexible process, namely the compression level varies within replayed episodes: Some fragments reappear on a timescale that resembles the original perception and replay is less compressed within sub-events of continuous episodes, which suggests that participants were able to flexibly skip between sub-events during memory replay.

## Acknowledgements

The authors would like to thank the Sir Peter Mansfield Imaging Centre (SPMIC), specifically Dr. George O’Neill, Dr. Benjamin A.E Hunt and Dr. Lauren Gascoyne for their help with data collection.

## Author Contributions

Conceived and designed the experiments: SM BPS SH.

Performed the experiments: SM.

Analyzed the data: SM under supervision of SH.

Contributed reagents/materials/analysis tools: SM SH HB.

Wrote the paper: SM SH commented and edited by BPS HB.

## Methods

### Participants

#### Balancing pilots

The balancing pilots were realized to balance the difficulty between videos that later served as scene 1, 2 and 3 of a video-episode. For each of the 2 balancing pilot experiments, 18 subjects were tested (36 total). In the first balancing pilot, 16 female and 2 male, right handed subjects participated that were on average 18.67 years old (youngest: 18, oldest: 20). In the second balancing pilot, 15 female and 3 male right handed subjects were tested. Their average age was 21.39 years (youngest: 18, oldest: 47). 2 additional subjects were tested in balancing pilot 2, however their behavioral performance was at chance and they were excluded from the analysis.

#### Behavioral pilot and experiment

For the behavioral pilot only 12 subjects (8 female, 4 male) participated that were on average 22.58 years old (youngest: 19, oldest: 29). 2 of the female participants were left handed, the rest were right handed. Data from 24 right handed volunteers (18 female, 6 male) was acquired for the behavioral experiment. The average age of this sample amounted to 22.79 years (youngest: 20, oldest 34).

#### MEG experiment

For the MEG experiment 24 volunteers (13 male, 11 female) participants were tested. Subjects were between 18 and 34 years old (mean: 23.92 years). 6 participants were left handed, 18 participants were right handed. 1 of the 24 subjects was excluded after pre-processing because of a persistent electrical artifact in the data that could not be removed with filtering.

6 additional subjects (4 female, 2 male) aged 19 to 28 years (mean: 22) were recorded but not analyzed. 2 subjects moved excessively throughout the recording session (maximal movement: 1.8 cm and 2.7 cm), 1 subject moved excessively throughout the session (maximal movement 1.4 cm) and fell asleep during the experiment. 1 subject felt unwell and aborted the experiment after approx. 10 % of the recording session, 1 subject only completed approx. 70 % of the recording session and moved more than 2 cm throughout the experiment. Finally 1 subject was lost due to technical failure during the recording. After preprocessing, the maximal movement of included participants across all trials (i.e. the range of all positions) was on average 5.89mm (s.d. = 2.62, min = 1.69, max = 9.09).

All included and excluded participants in the pilot studies, behavioral experiments and the MEG experiment, were native English speakers. Before participation they were screened for any neurological or psychiatric disorders. Their informed consent was obtained according to the ethical approval that was granted by the University of Birmingham Research Ethics Committee (ERN_15– 0335A), complying with the Declaration of Helsinki.

### Material and experimental set up

#### Videos

For each of the balancing pilots, a total of 12 short video-clips were used. Videos stemmed from a pool that was provided by Landesfilmdienst Baden-Württemberg, Germany, some of them were additionally edited. Each video-clip was a 2-second-long colored, dynamic scene that featured a single action (i.e. a ship sailing or a diver jumping into the water). During the task, video-clips were always superimposed with a transparent text box (white box with alpha value 0.9) in which the word-cue could appear. According to the behavioral results from balancing pilot 1, we edited or changed some of the scenes before the second balancing pilot. The final video-clips were 12 different scenes that belonged to four general topics. For the behavioral experiments and the MEG experiment, the video-clips were then grouped into four seamless sequences of frames that formed a video-episode (i.e. a sequence of three scenes that belong to a general topic and form a short story). The 3 scenes of each video-episode were clearly distinguishable.

According to the second balancing pilot, scenes that were assigned to be in 1st, 2nd or 3rd position of video-episodes, did not differ significantly in difficulty (percent correct responses), when associated with a word-cue. Pairwise comparisons with t-test of positions 1 and 2 (*t*_17_ = 0.86, *p* = 0.4), 2 and 3 (*t*_17_ = 0.15, *p* = 0.88) and 1 and 3 (*t*_17_ = 1.4693, *p* = 0.16) and Bayes-Factor analysis supported the null Hypothesis of no difference between positions. This was supported either by substantial (*BF*_01_>1.6) or strong (*BF*_01_>3.3) evidence for the comparison of positions 1 and 2 (*BF*_01_ = 2.97) of positions 2 and 3 (*BF*_01_ = 4.07) and of positions 1 and 3 (*BF*_01_ = 1.65). Importantly, reaction times in the second balancing pilot did not differ significantly between the video-clips that we finally assigned to be in position 1, 2 or 3. Pairwise comparisons with t-test of assigned positions 1 and 2 (*t*_17_ = −0.59, *p* = 0.56), 2 and 3 (*t*_17_ = −0.31, *p* = 0.76), and 1 and 3 (*t*_17_ = −1, *p* = 0.33) and Bayes-Factor analysis supported the null Hypothesis for the comparison of positions 1 and 2 (*BF*_01_ = 3.53) of positions 2 and 3 (*BF*_01_ = 3.95), and positions 1 and 3 (*BF*_01_ = 2.67).

#### Word-cues

Word-cues were downloaded from the MRC Psycholinguistic Database ^30^. For the balancing pilots, we divided 540 word-cues into 18 lists. Those lists did not differ in Kucera-Francis written frequency (mean = 20.80, s.d. = 8.55), concreteness (mean = 506.50, s.d. = 90.07), imageability (mean = 521.04, s.d. = 69.51), number of syllables (mean = 1.63, s.d. = 0.68), number of letters (mean = 5.61, s.d. = 1.42) or word-frequencies taken from SUBTLEXus (mean = 15.22, s.d. = 14.07); specifically, “Subtlwf” was used ^31^. In the balancing pilots, 12 of the lists were associated with a video-clip and 6 of the lists were assigned to become a distractor word. Across subjects each list was associated with every movie once and served as a distractor word six times. This was done to additionally control for list specific effects across subjects. An additional 9 words were randomly selected for practice.

For the behavioral pilot, the behavioral experiment and the MEG experiment, we divided 360 word-cues into 12 lists. Those lists were likewise balanced for Kucera-Francis written frequency (mean = 20.41, s.d. = 7.47), concreteness (mean = 518.72, s.d. = 78.39), imageability (mean = 530.78, s.d. = 60.17), number of syllables (mean = 1.56, s.d. = 0.62), number of letters (mean = 5.44, s.d. = 1.30) and word-frequencies taken from SUBTLEXus (mean = 15.07, s.d. = 13.04); again, “Subtlwf” was used ^31^. Across participants, each of the lists was associated with every video-clip twice. An additional 6 words were randomly selected for practice.

#### Response options

To create the response options (see Fig. 1c-d), we took Screenshots from the video-clips. In the balancing pilots, we adjusted brightness and contrast, so that no screenshot appeared more salient. For the behavioral pilot, the behavioral experiment and the MEG experiment the numbers 1, 2 and 3 were framed by a square which resembled a frame from an old film. Those represented the first response options, i.e. the choice between scene 1, scene 2 or scene 3. For the second response, i.e. the response about the correct video-episode, the 3 screenshots from the concatenated video-clips were presented next to each other for each of the 4 choices. In the control condition of the behavioral experiment, the response option intact/rearranged was realized with a screenshot which was of the same size as the videos during presentations. This screenshot was superimposed by a transparent textbox containing a word-cue. The words intact and rearranged were displayed at the left and right of the textbox as response options. The left/right position of these options was balanced across participants.

#### Behavioral setup

Visual content was presented on an LED monitor (Samsung syncmaster 940n at a distance of approximately 60 cm from the subject’s eyes. The monitor was set to a refresh rate of 60 Hz. On a screen size of 1280 x 1024 pixels, the video-clips had the dimension of 360 pixels in width and 288 pixels in height on the screen. “Helvetica” was chosen as the general text font, font size was set to 22 for instructions and to 28 for word-cues. Black text (rgb: 0, 0, 0) and movies were presented against a white background (rgb: 255, 255, 255).

#### MEG setup

MEG was recorded at the Sir Peter Mansfield Imaging Center (SPMIC) in Nottingham, UK. Subjects performed the experiment in a seated position at a distance of approximately 60 cm from a white screen. The image was projected onto the screen using a PROPixx projector (VPixx Technologies, Saint-Bruno, Canada) that operated at a refresh rate of 60Hz and a resolution of 1920 × 1080 px. The projected image appeared at a size of approx. 40 × 22.5 cm on the screen. Accordingly, the video-clip appeared in a dimension of approx. 15 × 12 cm. An eye tracker (EyeLink 1000 plus, SR Research, Ontario, Canada) was placed in front of the screen. The tracker was mounted in an upwards facing orientation, slightly below the visible display, on a small wooden board. In this setup it tracked the subject’s left eye from below and from a distance of approximately 55 cm.

### Procedure

#### Balancing pilots

The balancing pilots were realized to ensure that no material specific differences between the first, second or third position of a video-episode were to be expected in the following experiments. To this end, the video-clips that were considered were presented as single scenes during learning blocks, where they were superimposed by a transparent text box, containing a word-cue. Upon informed consent and completion of screening questionnaires, participants sat down in front of the screen and received a standardized instruction for the task. All subjects saw the video-clips for familiarization and completed a practice version of the task before starting. Participants performed 15 runs of an encoding block, in which their task was to vividly associate the word-cue with the corresponding video-clip, a short distractor block, in which they did some easy math and a retrieval block in which they retrieved the associated video-clips as quickly as possible, whilst presented with a word-cue.

During encoding a fixation cross was displayed for 2 seconds. Then the video-clip and word-cue played for 2 seconds. Finally, a fixation cross appeared again for 2 seconds. In every encoding block, each video-clip was presented twice amounting to a total of 24 trials in every block. The video-clips were presented in a balanced but randomized order such that no movie was presented more than 2 times in a row.

In the distractor block, subjects solved simple math problems. For 45 seconds they were either presented with the word bigger or with the word smaller and two single digit sums (e.g. 4+5 and 3+2). Their task was to select the correct sum (i.e. either the bigger or the smaller sum). Feedback was given in the form of the words “correct” and “wrong” appearing in green and red respectively on the screen.

For each retrieval block the current 24 cues were mixed with 12 new distractor words in a randomized way, such that items corresponding to the same video-clip or items corresponding to a distractor did not appear more than 2 times in a row.

In the retrieval block subjects were asked to select the video corresponding to the word-cue as fast as possible. The target (i.e. the screenshot from the correct clip) and two lures (two screenshots from a different clip) were presented on three positions around the word-cue. The positions formed a triangle with equal distance from the center to the left and right and 2/3 of that distance above the word-cue. In addition to those three response options, a question mark was displayed below the center of the screen. This response option was available in order to indicate that no video-clip-screenshot was identified as the correct target.

In order to control for effects from specific screen positions, the mapping of targets to positions was randomized but balanced, such that the target was presented on every position 8 times. To control for item specific effects, the two lures that were presented with the target were assigned, such that every video-clip-screenshot served four times as a lure to a target and the same screenshot was never on both lure-positions. The 12 additional distractor words were, by definition, only paired with lures, we balanced the random mapping, such that every video-clip-screenshot served once as a lure on every position.

In order to respond, participants placed the index finger of their dominant hand on the number 2 of the numeric keypad of the keyboard. The index finger rested there as long as no response was required. When presented with the cue-word subjects could either press one of the numbers 4, 6 and 8 which corresponded spatially to the response options (screenshots) on the screen and were in approximately the same distance from the starting position (number 2), or they could press 0 which corresponded to the question mark. Available buttons were highlighted with colored stickers to facilitate orientation.

Whilst presented with the word-cue, subjects had maximally 4 seconds to select their answer. At the end of every retrieval block participants were reminded that associations from the previous block were now irrelevant and had the opportunity to take a self-paced break.

#### Behavioral pilot and behavioral experiment

In the behavioral pilot (Fig. 1a, b, d top-left), subjects saw video-episodes that consisted of 3 distinct scenes. Those scenes comprised of the video-clips from balancing pilot 2, which ensured that no material specific differences were to be expected between position 1, 2 and 3 of the video-episodes; not in memory performance and most importantly not in reaction time. Participants first completed the screening questionnaire and gave informed consent. After instruction with the task, they saw the video-episodes twice for familiarization and were instructed to pay attention to their 3-scene-structure, such that they could confidently identify the first, second and third scene of each video-episode.

After a short practice version of the task, the experiment started. It was again a sequence of encoding, distractor and retrieval blocks. In each encoding block subjects learned a series of associations (Fig. 1a). They first saw a fixation cross on the screen for 2 seconds. After that one of the four video-episodes played for 6 seconds. During this video-episode a transparent textbox was overlaid on the video. In one of the three scenes, a word-cue appeared in the textbox and disappeared again with the end of the scene. Subjects were instructed to form a vivid association between the word and the precise scene of the video-episode, such that they could later recall that exact scene and video-episode upon presentation with the word-cue. We randomized the presentation of the associations in a balanced way, such that no video-episode was presented more than twice in a row and a word-cue did not appear in the same position more than twice in a row. Additionally, every position within every video-episode was associated with a word cue once within 12 subsequent associations.

After each video-episode a fixation-cross showed for 1 second then subjects rated the plausibility of the association between word-cue and scene. Three response options were labelled with “not plausible”, “plausible” and “very plausible” and could be selected with the buttons 4, 5 and 6 on the numerical pad of the keyboard. The plausibility rating served to keep participants engaged in the task and support memory formation. In the distractor block, subjects were presented again for 45 seconds with simple math problems and had to decide which one of two single digit sums was either bigger or smaller (Fig. 1b). For the retrieval block the word-cues were now randomized again in a balanced way, such that word-cues corresponding to the same video-episode regardless of position, or to the same position regardless of video-episode, did not appear more than twice in a row.

Retrieval blocks (Fig. 1d, top-left) started with a fixation cross, displayed for 2 seconds. Then a word-cue appeared in the center of the screen and the three framed numbers appeared on a triangle around the word-cue. Participants were instructed to select, as quickly as possible, in which of the three scenes they learned the word. For this choice they only saw the numbers 1, 2 and 3; after they made this choice, screenshots forming the four video episodes appeared in the four corners of the screen. Participants were asked to indicate now, to which of the four episodes the selected scene belonged. The position of the numbers 1, 2 and 3 as well as the mappings of the four screenshot-sequences to the screen positions were randomized in a balanced way, namely all possible permutations of 1, 2 and 3 were randomly mapped onto the three positions within 6 subsequent trials and all possible permutations of the four positions of the video-episode screenshots were used within 24 trials. This was done to control for any potential effects from specific screen positions on reaction times or position specific response preferences. In order to respond, volunteers were asked to place the index finger of their dominant hand on the number 5 of the numerical pad on the keyboard. The surrounding numbers 4, 6 and 8, which form a triangle around the number 5 were highlighted with red stickers and served as the response options for the scene-response (first response: 1, 2 or 3). Those buttons corresponded spatially to the position of the permuted numbers 1, 2 and 3 on the screen. Accordingly the buttons 1, 7, 9 and 3 which form a square on the numerical pad, were available for the second response which informed about the correct video-episode. Importantly subjects were instructed to make all responses with the index finger of their dominant hand and go back to the starting position after every response, i.e. leave the finger resting on the button 5. At the end of every retrieval trial, a scale appeared on which subjects rated the confidence in their response. Three options were labelled with “guess”, “sure” and “very sure” and corresponded to the buttons 4, 5 and 6 on the numerical pad.

Participants performed a variable amount of runs of encoding, distractor and retrieval blocks that varied in length according to their individual memory performance. The first block comprised of 24 items, subsequently its length was adjusted. If more than 70% of items were recalled correctly in the last block (i.e. correct scene and movie were selected), 12 items were added to the next block, if less than 50% were recalled correctly, 12 items were removed from the following block. All blocks comprised at least of 12 associations that had to be learned. All participants completed 360 trials in total.

In the final behavioral experiment (Fig. 1a, b, d top row) subjects performed exactly the same task as in the behavioral pilot experiment, however, every other block was performed with a different retrieval task. Specifically subjects performed the same learning paradigm, yet they did alternating retrieval blocks of cued-recall (CR, see above) and associative recognition (AR). In the AR blocks (Fig. 1d, top-right) subjects were presented with a screenshot of a single video-clip, representing one of three scenes within a video-episode. The center of the screenshot was again superimposed with the transparent textbox containing one of the previously learned word-cues. The association between word-cue and video-clip could either be intact, i.e. the word was learned in this exact position within the video-episode, or it could be rearranged. In the latter scenario, a different video-clip from the same video-episode was superimposed by the word-cue. This means that word-cues were either presented in the correct position or in the wrong position within the video-clip. Participants were again instructed to decide as quickly as they could, whether the association was intact or rearranged. Block-size was adjusted in the same way with percent of correct responses measured as 200*(Hits - False Alarms)/N, with Hits being the number of correctly identified intact associations and False Alarms referring to the number of rearranged associations that were declared intact and N referring to the number of trials in the last block. Response buttons for the intact/rearranged choice were 4 and 6 on the numerical pad, which are in equal distance from the number 5, where the index finger of participants’ dominant hand rested comfortably at the beginning of each trial. After the experiment participants answered a few interview questions regarding eventual strategies and their subjective experience of the task.

#### MEG experiment

In the MEG experiment (Fig. 1a, b, c) volunteers learned associations between video-episodes and word-cues in the same way as in the behavioral experiment. Memory retrieval was similar to the behavioral pilot experiment (i.e. a cued-recall task); however a fast response was not required (see below). Upon informed consent and screening questionnaires, participants received the instructions for the task on a laptop outside the scanner. They familiarized themselves with the video-episodes twice, paying close attention to their structure. It was ensured that every participant was able to identify the three different scenes of a video-episode. In a short practice, they performed a block of encoding, distractor and retrieval with the six example words. The head-localization coils of the MEG system were attached to the participants’ head and their positions were logged along with the shape of the participant’s head (see Data Collection). Subsequently, volunteers were seated in a comfortable position under the MEG helmet. Subjects used a single button on each of two response pads with their left and right index finger. After the eye tracker was mounted and calibrated, the experiment started.

The MEG experiment was again a sequence of encoding, distractor and retrieval blocks. In each encoding block (Fig. 1a), subjects learned a series of associations between scenes in video-episodes and unique word-cues. Participants first saw a fixation-cross on the screen for 1 second. After that one of the four video-episodes played for 6 seconds overlaid with a transparent textbox. In one of the three scenes of the video-episode, the unique word-cue appeared in the textbox and disappeared again with the end of the scene. The task was again to form a vivid association between the word and the precise scene of the video-episode in order to later recall the exact scene and video-episode, when only presented with the word-cue.

After the video-episode, the fixation-cross appeared again for 500ms. Finally, the two response options ‘plausible’ and ‘not plausible’ appeared on the left and right of the screen. Subjects used the left or right button to indicate whether the association between video-scene and word-cue was plausible to them. This task kept participants engaged and supported their memory performance.

The order of presentation was randomized in a balanced way: no video-episode was presented more than twice in a row and a word-cue did not appear in the same position more than twice. Additionally every position within every video-episode was associated with a word cue once within 12 subsequent associations. In the distractor block (Fig. 1b) subjects solved simple math problems for 45 seconds: They had to decide which one of two single digit sums was either bigger or smaller, using a left or right button press. Participants received feedback on the distractor task in form of the words “correct” or “wrong” displayed in green or red respectively. For the retrieval block the word-cues were now randomized again in a balanced way, such that word-cues corresponding to the same video-episode regardless of position, or to the same position regardless of video-episode, did not appear more than twice in a row.

Trials of the retrieval block (Fig. 1c) started with a fixation cross that was displayed for 1 second. Then a word-cue appeared in the center of the screen for 3.5 seconds. In this time interval subjects remembered in which exact scene they had seen this word. For a random time interval between 250ms and 750ms, a fixation cross was shown again, then the response options appeared. The time interval for retrieval was chosen based on reaction-time data from the behavioral experiments, such that participants could comfortably remember the correct association. The first response option required the selection of the correct scene. To this end pictograms featuring the numbers 1, 2 and 3 were displayed on the top of the screen. The mapping of the numbers 1, 2 and 3 to the three screen-positions was randomized in a balanced way such that all possible permutations appeared within 6 subsequent trials. Participants could now move a red square, which framed the current selection. By pressing the left button they changed their selection by moving the frame clockwise. This selection was confirmed by pressing the right button. Note that this button assignment ensured that subjects would always prepare the same response during the retrieval trial, regardless of the memory content. This is important to control for trivial but systematic differences that correlate with memory content in the retrieval interval.

After the position was selected, the two other position pictograms were overlaid with transparency (alpha = 0.9), such that the selected option remained highlighted on the screen. The concatenated screenshots from the video-episodes appeared below the position-pictograms and the red selection frame could be moved clockwise with the left button. Again the selection was confirmed with the right button. To ensure that subjects followed instructions and tried to recall the correct position as soon as they were presented with the word-cue (and did not wait until the response options were presented), there was a time limit of 4 seconds to select the correct position and again to select the correct movie. To allow for flexibility due to hasty or imprecise selections, 200ms were added to this time limit, whenever the selection-frame was moved. Participants did not know about this increment; all participants selected their responses quickly but not hastily. If the time limit was exceeded, the message ‘too slow’ appeared at the center of the screen for 5 seconds. Altogether the time limits were designed, such that subjects could comfortably remember the correct association during the presentation of the word-cue, and were eager to select the two responses straight away. After the associated video-episode was selected, unselected response options were overlaid with transparency for 300ms, then the two options ‘guess’ and ‘know’ were presented on a new screen to give the participant the opportunity to communicate, whether the selected answers were based on a guess.

Participants performed a variable amount of runs of encoding, distractor and retrieval blocks. The blocks varied in length according to individual memory performance. The first block comprised of 24 items, subsequently its length was adjusted. If more than 90% of items were recalled correctly in the last block (i.e. correct scene and movie were selected), 24 items were added to the next block, If more than 70% of items were recalled correctly, 12 items were added to the next block, if less than 50% were recalled correctly, 12 items were removed from the following block, if less than 40% were recalled correctly, 24 items were removed. All blocks comprised at least 12 associations that had to be learned, i.e. block-size was never reduced below 12 items; all participants learned and recalled a total of 360 associations.

### Data Collection

Stimulus presentation and the collection of behavioral data was realized on a standard desktop computer running MATLAB 2014b (MathWorks) under Windows 7, 64 Bit version. Stimuli were presented through the Psychophysics Toolbox Version 3 ^32^. In the behavioral experiments, responses were collected from button presses on the numerical pad of a wired keyboard (Model 1576, Microsoft Corporation, Redmond, US). In the MEG experiment, fiber optic response pads were used.

Neurophysiological data were collected with 275-channel CTF MEG (CTF, Coquitlam, BC, Canada) at the Sir Peter Mansfield Imaging Center (SPMIC) in Nottingham, UK. The system was used in third-order gradiometer configuration, recording at a sampling frequency of 600 Hz over the whole duration of the experiment. Three localization coils that were attached to the participants’ left preauricular point (LPA), right preauricular point (RPA) and to a point slightly above the nasion (NAS) were energized during the recording session. This was done to localize the head position relative to the sensors.

Head digitization was collected with a Polhemus ISOTRAK device (Colchester, Vermont, USA). A minimum of 500 points on the scalp were logged relative to the positions of the three fiducial points (LPA, RPA, NAS). Individual anatomical data was acquired via magnetic resonance imaging (MRI) (3T Achieva scanner; Philips, Eindhoven, the Netherlands) with an MPRAGE sequence covering the whole head at 1mm^3^ resolution. MRIs were either measured at the SPMIC or at the Birmingham University Imaging Centre (BUIC).

For 17 of the included subjects (23), eye tracking (Eyelink 1000 Plus, SR Research, Ontario, Canada) was recorded on a separate Computer provided by the manufacturer at a sampling rate of 2000 Hz. The data was additionally written into 3 analog input channels of the MEG system via the EyeLink Analog Card. The eye tracker was used in remote mode tracking the pupil and corneal reflection with a 16mm lens. It was calibrated and validated using 13 points on 80% of the screen, which contained all of the task relevant information.

### Analysis of Reaction Times

We defined reaction time (RT) as the time to the first response after onset of the word-cue (Fig. 1d, bottom-row). All RTs faster than 200ms were considered implausible and discarded from further analysis. Additionally RTs that were 2.5 standard deviations above the mean RT were discarded. The means of remaining RTs were then tested statistically. To account for the non-normal distribution of RTs ^33^, all statistical tests are also reported for log-transformed RTs.

### Preprocessing of Neural Data

The data was preprocessed in MATLAB 2015a (MathWorks) with a combination of functions from the Fieldtrip toolbox for EEG/MEG analysis ^34^ and custom written scripts.

For the sensor level analysis, the 3^rd^ order gradiometer correction was first applied, then the continuous recording was filtered with a Butterworth IIR filter of 4^th^ order with a stopband of 49.5 to 50.5 and its harmonics (99.5 - 100.5, 149.5 - 150.5, 199.5 - 200.5, and 249.5 - 250.5) to reduce the line noise artifact. Additionally the data was filtered with a stopband of 59 – 60 to attenuate noise with a center frequency of 59.5 Hz.

Subsequently, the data was segmented into trials that started 1.5 seconds prior to video-onset and ended 7.5 seconds after video-onset at encoding. Trials at retrieval started 1.5 seconds prior to the onset of the word-cue and ended 5 seconds after onset of the word-cue. If available, the dataset was combined with the downsampled and segmented trials from the eye tracking.

To remove activity from eye blinks and noise, and to detect heartbeats, Independent Component Analysis (ICA) was used ^35^. For the computation of the ICA unmixing matrix, trials containing coarse artifacts or showing strong muscle activity were heuristically excluded. Additionally, the data was downsampled to 250 Hz and cut to 1 second-long segments; the obtained unimxing matrix was then applied to all original trials.

When possible, we compared independent components with the eye tracking data; we removed those components that picked up eye-blinks or eye-movement related activity. Additional components that picked up electrical noise were removed from the data. A copy of components which contained a clear R-wave of the QRS complex in a heartbeat was stored for later peak-detection and regression. All remaining components were projected back to a channel representation.

Finally, all data was inspected visually and trials containing artifacts were removed from later analysis.

After visual inspection, 84.26 % (S.D. = 8.29 %) of trials remained.

Heartbeats were removed with a regression based approach: An iterative peak detection algorithm was applied to the ICA-component showing the clearest R-wave; it served as a proxy for ECG. This was done only for the remaining trials after visual inspection. Before peak-detection the heartbeat-component was highpass-filtered (4Hz, 4^th^ order Butterworth). The peak detection algorithm first calculated a plausible maximum of heartbeats that were not to be exceeded. The signal was z-scored and thresholded. Local peaks were detected by finding local maxima in clusters of z-scores that were above threshold. Subsequently the threshold was lowered stepwise, down to a z-score of 2. With lowering threshold, increasingly bigger areas around the peaks were excluded from further peak detection. If the maximum number of plausible peaks was exceeded, the threshold was no longer lowered. A heartbeat template was now created by averaging 500ms long segments around the peaks. Gaps in the continuous recording were subsequently zero-padded in order to convolve the component with the template. Peak detection was then repeated on the convolved time course and a new template was built from these peaks for subsequent convolution ^36^. After a few repetitions, the template converged and the resulting peaks were controlled manually, even though errors rarely needed to be corrected.

Instead of simply subtracting the averaged template from the data, the trials were now split into four big segments and a general linear model (GLM) was built around the peaks in each segment. A high pass filter (1Hz, 4^th^ order Butterworth) was applied to the data, only for the purpose of fitting the model. The GLM consisted of a separate repeated measure factor for each time point in the heartbeat, beginning 280ms before the peak and ending 720ms after the peak ^36^. Additionally, a separate factor was included for every heartbeat, which modelled the offset between 280ms pre-peak and 720ms post-peak. Furthermore an offset factor for the overall segment was included. The solved model was then applied to every channel. The data model ŷ was built by using only the repeated measure factors, which modelled each time point within the heartbeat (i.e. the beta weights for offsets were set to 0). After visual inspection, this resulting model of the heartbeat was subtracted from each original channel.

For the source level analysis, the anatomical data was first aligned to the digitized head positions. This was done by extracting the surface of the head from the anatomical MRI; in a first step a rough alignment was done manually, then the Iterative Closest Point (ICP) algorithm implemented in fieldtrip ^34^ was used to match the surface to the point-cloud of the head digitization, finally this solution was controlled and eventually corrected again manually. The transformation to the aligned space was subsequently applied to the segmentation of the brain, which was likewise extracted from the anatomical images. To correct for head movements, the average head positions within the trials were first clustered, such that one positional-cluster was built for every 10 trials. Subsequently a separate lead field was computed for every cluster and then averaged. Hereby, an average lead field across all trials was obtained for each participant ^37^. Importantly ‘all trials’ refers to the trials that were included in a given contrast (e.g. for the contrast of Hits and Misses at retrieval, encoding trials were not included in the computation of the lead field). Before the source level analysis, the 3^rd^ order gradiometer correction was applied to the cut raw-data, lead fields were adjusted accordingly. Finally the data was demeaned and bandpass filtered between 4 and 15 Hz. The position of virtual sensors in individual brains was derived from a 1 cm spaced grid, which was placed 6mm below the surface of the cortex into the MNI brain and then spatially warped into individual brains. This was done via the inverse of the transformation describing their normalization and resulted in 1407 individual virtual sensor positions which were anatomically equivalent. Finally, to reconstruct activity on virtual sensors a linearly constrained minimum variance (lcmv) beamforming approach, implemented in the Fieldtrip toolbox ^34^, was used. Filter coefficients were again computed on all data in a given contrast.

### Analysis of oscillatory power (Supplemental Information)

To estimate oscillatory power at retrieval (Supplemental Fig. 1), the Fourier-transformed data was multiplied with a complex Morlet wavelet of six cycles. This was done in steps of 10ms for every full frequency between 2 and 40Hz. The raw power was then obtained from the squared amplitude of the Fourier spectrum. Across all trials within the contrast (i.e. Hits and Misses), a baseline was computed as the average power between 1 second pre-stimulus and 4 second after stimulus onset ^38^. Trials were then normalized by subtracting the baseline and dividing by it (activity_tf_ - baseline_f_)/baseline_f_, with t indexing time and f indexing frequency.

### Region of Interest (ROI)

An occipito-parietal region of interest (ROI) was derived from the AAL atlas ^39^ (Fig. 2b). To obtain the ROI in form of a group of virtual sensors, the sensor-positions in MNI-space were assigned to the nearest described AAL-region, based on their Euclidean distance. The occipito-parietal ROI comprised of bilateral AAL-regions: angular gyrus, calcarine sulcus, cuneus, inferior occipital cortex, inferior parietal lobule, lingual gyrus, middle occipital gyrus, precuneus, superior occipital gyrus, superior parietal lobule, supramarginal gyrus.

### Content specific oscillatory phase at encoding

During encoding, participants repeatedly watched the same video-episodes. Hence, it was possible to assess content specific properties if they were more similar between trials of same content than between trials of different content (Fig. 2a). In order to determine whether the ongoing oscillatory phase was specific to individual perceptual content, trials were grouped into 4 sets according to the video-episode that was perceived. The complex Fourier spectrum was again derived by multiplying the Fourier-transformed data with a complex Morlet wavelet of six cycles. Then, inter-trial phase coherence ^40^ (ITPC) was computed across the trials of same content (i.e. for each of the four trial-groups). This was done at every full frequency between 2 and 40 Hz in steps of 10ms starting 1 second before the onset of the video-episodes and ending 7 seconds after the offset of the video-episodes. Following that, the trials were shuffled and grouped randomly into 4 sets of mixed-content-trials. Sets were of equal size to the 4 sets of same-content-trials. Again ITPC was computed separately for each of the 4 sets. To balance the contribution of the 4 sets, a Rayleigh Z-correction was applied with N*ITPC^2^, where N refers to the number of trials in a set. Finally the corrected ITPC was averaged across the 4 sets in the ordered and in the shuffled condition. Their difference indicated content specificity of phase which could be statistically tested ^16,41^. The analysis in source-space was done in the same way using the virtual sensors; however the frequency was restricted to 8 Hz.

### Content specific phase similarity between encoding and retrieval

The reactivation of temporal patterns (Fig. 2c-d, Supplemental Fig. 3) was estimated on virtual sensors for the frequency of 8 Hz. To this end, the oscillatory phase coherence between encoding and retrieval was contrasted between trial-combinations of same content (e.g. watching video-episode A, recalling video-episode A) and random trial-combinations of different content (e.g. watching video-episode A, recalling video-episode B). The combinations were balanced, such that in both conditions (same vs. different combinations) exactly the same trials were used in the same amount of combinations. We only changed the pairing between encoding and retrieval trials. For each trial-combination, 1-second long windows from the encoding trial were now compared to every time point at retrieval starting at the onset of the word-cue and ending at its offset after 3.5 seconds. This comparison was done with a sliding window approach. As a metric of phase-similarity, the phase coherence across time ^8,18,19^ (i.e. across the 1 second window) was computed. All possible windows from encoding were used in this sliding window approach, with the first window ranging from 0 to 1 seconds and the last window ranging from 5 to 6 seconds during the video-episode (compare Fig. 2 c). Note that the response options set on between 250ms and 750ms after the word-offset, additionally the first response-screen did not contain content-information (only the numbers 1, 2, and 3) and all responses required a button-press on the left button. Therefore, no confounds from the response interval were expected to bleed into the tested interval. Oscillatory phase was estimated by multiplying the Fourier-transformed data with a complex Morlet wavelet of six cycles in steps of 15.6ms for consistency with our previous analyses ^8^. The average similarity between all time-windows and combinations was subsequently averaged to derive a single value of similarity for combinations of same content and a single value for combinations of different content at each virtual sensor. Note that this method enables the investigation of highly dynamic patterns in a robust way, because a measure that captures dynamic changes in ongoing oscillations is accumulated across encoding time, retrieval time and ten thousands of trial-combinations.

### Time courses of Replay

To observe the temporal scale of reactivation (Fig. 3), the distribution of similarity to the remembered stimulus content (i.e. phase coherence) across retrieval was compared between different sliding windows from encoding. By definition, a distribution is normalized to an area under curve of 1 and therefore accounts for differences in total similarity between windows. To robustly compare the distribution of similarity between 6 non-overlapping windows, phase-coherence was accumulated across time, such that at the beginning of the retrieval time, zero similarity to all windows was present and at the end of retrieval (i.e. at 3.5 seconds after word onset) 100 % of similarity was reached (Fig. 3c). This made it possible to compare at each time point, whether the similarity to a window had come up earlier than to another window. In other words: If patterns from window “A” tend to appear earlier than patterns from window “B” across subjects, then the cumulated similarity to window A should be statistically higher than the cumulated similarity to window “B”, at several time points.

In order to test for a general tendency for forward replay, a line was fitted across all 6 windows and tested against a slope of 0 (Fig. 3d). Hence a negative slope of this line means that earlier windows from encoding appear earlier during retrieval. In order to test the hypothesis that the replay of individual scenes takes place on a slower timescale (Fig. 3e), 3 lines were fitted across the 2 non-overlapping windows within each scene, and their slope was averaged. A more negative average slope of these 3 lines compared the slope of the line across all windows supports the hypothesis that replaying individual scenes takes place on a slower temporal scale.

Importantly this way of cumulating the similarity distributions allows for robust testing across subjects at the expense of introducing temporal dependencies between time points. Specifically, if more similarity to a window is present at an early point this can propagate to later points, if similarity thereafter increases at the same speed for all windows. The extent of significant time intervals should therefore be interpreted with caution. Another disadvantage of this method is that the slope is interval scaled and its absolute value is not interpretable.

In order to compensate for this disadvantage and quantify the actual lag between time windows from encoding descriptively (Fig. 3a-b), the distributions of similarity were averaged across subjects and smoothed with a moving average kernel of 250ms, to attenuate noise (Fig. 3a, right). The cross-correlation between distributions was then computed to estimate the lag between them: The shape of one similarity distribution is matched to another (Fig. 3b). This was done within the time interval in which the slowing down of replay was observed; specifically in which the slope for lines fitted within a scene was significantly more negative than the slope across all windows (i.e. between 550ms and 2350ms at retrieval).

### Statistical analyses

#### Behavioral performance and Reaction times

Behavioral performance was tested with a repeated-measures-ANOVA, on the percent of correct responses. Post-hoc tests were then performed with 2 separate ANOVAs for the final behavioral experiment and with a series of one-sample t-test (see Supplemental Information).

RTs in the balancing pilots were first contrasted with one-sample t-tests. In order to statistically test the null hypothesis the Scaled JZS Bayes Factor ^42^ to the one-sample t-tests was computed. RTs in the behavioral pilot experiment were compared with a repeated-measures-ANOVA with the factor position (1, 2 and 3). In the final behavioral experiment, a 2×3-repeated-measures-ANOVA was computed with the factors retrieval task (cued-recall vs. associative recognition) and position (1, 2 and 3). Post-hoc tests were then performed with 2 separate ANNOVAs. Reaction times for the 3 different positions were subsequently compared with a series of post-hoc one-sample t-tests. Greenhouse-Geisser correction was used with all ANOVAs, null-effects of interest were tested with Bayesian t-tests ^42^.

#### Content specific oscillatory phase at encoding

Content specific phase at encoding was statistically tested by contrasting average ITPC across arranged groups with the average ITPC across shuffled groups. This was done with a series of t-test at every time point between 0 and 6 seconds after onset of the video-episode, at every frequency between 2 and 40 Hz and at every sensor. Multiple comparison correction was done via Monte-Carlo permutation of contrast labels as implemented in the fieldtrip toolbox ^34,43^. 3-dimensional clusters and cluster-sums were formed across time, frequency and sensors. Cluster-sums in the original contrast were compared to the distribution of cluster-sums under random label assignment in order to derive p-values. The cluster-forming threshold corresponded to the critical t-value (alpha < 0.05) of a single-sided one-sample t-test, 1000 random permutations were drawn. On the source level content specific phase was assessed for the frequency of 8Hz. Again the ITPC of arranged groups and the ITPC of shuffled groups were contrasted with a one sample t-test that was computed at every time point and every virtual sensor. Clusters were summed across neighboring sensors and time points in 1000 random permutations. To obtain time courses within the parieto-occipital ROI, t-values were averaged across all virtual sensors within the ROI.

#### Content specific phase similarity between encoding and retrieval

Based on previous results ^8^, statistical testing for content specific reactivation was done for the frequency of 8 Hz, restricted to an occipito-parietal region of interest (ROI) derived from the AAL atlas ^39^. Averaged similarity values of encoding-retrieval combinations were contrasted between combinations of same content and combinations of different content. This was done with a one-sample t-test on every virtual sensor within the ROI. Subsequently t-values were thresholded with a t-value corresponding to a one-sided alpha value of 0.05; clusters were built across neighboring virtual sensors. Statistical testing was done again via 1000 random permutations. Cluster-sums in the original contrast were compared to the distribution of cluster-sums under random label assignment in order to derive p-values. A series of post-hoc t-tests was done on every time-point at retrieval in order to estimate the contribution to the effect from encoding windows (see Supplemental Information, specifically supplemental Fig. 3a).

#### Time courses of replay

Time courses were obtained by averaging across the ROI, which allows for an unbiased investigation of the time-courses of reactivation (see Supplemental Information, specifically supplemental Fig. 2a). Specifically, the cluster correction approach results in a biased noise-distribution within the cluster of significant reactivation. This renders the interpretation of its shape and any post-hoc analysis on sensors within the cluster problematic ^43^, see also ^44^. Since 86.46% of the t-values in the ROI were positive, we therefore decided to average across all virtual sensors within the anatomical ROI for the analyses of all time courses that were statistically tested.

Likewise, similarity densities were computed on the averaged similarity values across all virtual sensors within the ROI. The cumulated similarity density distributions for 6 non-overlapping encoding-windows were obtained for every subject. Consequently at every retrieval time-point a line could be fitted across 6 values for every subject. The slope of that line was subsequently subjected to a t-test against 0 across all subjects. The resulting time-course of t-values across the whole retrieval time was finally subjected to a multiple comparisons correction by controlling the false discovery rate ^45^. To compare the speed of replay within scenes, to the overall speed, the average slope fitted across two windows each (windows within scenes) was statistically tested against the slope across all encoding windows with a series of one-sample t-tests. T-values were obtained again at every time point during retrieval and the false discovery rate was controlled in order to correct for multiple comparisons. To estimate at which time-points reinstatement could be detected best (Supplemental Information, Fig. 2a), a series of one-sample t-tests was computed at every retrieval time point, between encoding-retrieval similarity of same content combinations and encoding-retrieval combinations of different content combinations (see Supplemental Information). Finally, the average similarity to all encoding time points was compared within the ROI, between trials in which an association from the first, second or third scene was recalled (Supplemental Information, Fig. 2b). This was done with a repeated-measures-ANOVA with the factor position and pairwise post-hoc t-tests.

#### Oscillatory power (Supplemental Information)

Baseline corrected oscillatory power was contrasted on the sensor level with a series of one-sample t-tests. Multiple-comparison correction was realized with a cluster-based Monte-Carlo permutation as implemented in the fieldtrip toolbox ^34^. 1000 permutations of contrast-labels were used; the clusters were formed from neighboring values below a threshold (see below). Neighboring values were derived across time from 0 to 4 seconds after the onset of the word-cue, across frequency from 2 to 40 Hz and spatially across sensors The threshold was the t-value which corresponds to a threshold of alpha = 0.05 for a single sided test. The maximal cluster-sum of real data was then compared to the distribution of maximal cluster-sums under random permutations in order to derive a p-value. In order to find the most robust frequencies that showed oscillatory power decreases, a t-test was computed for the average power difference across time (0 – 4s), sensors and frequencies. On the source level, baseline-corrected power at 8 Hz was averaged over time between 0 and 4 seconds and subjected to a one-sample t-test. Multiple comparison correction was addressed with the same cluster-based permutation approach; however, clusters were formed across neighboring virtual sensors.

## Supplemental Information

### Behavioral performance

In the behavioral pilot experiment, participants recalled the correct position and video-episode in 72.29% (*SD* = 11.69%) of the trials. A main effect of position indicated decreasing performance when an association had been learned in a later scene of a video-episode (ANOVA: *F*_1.27, 13.95_ = 4.988, *p* = 0.036, means: 74.86%, 72.29%, 69.72%); post-hoc tests indicated that only associations from the first position were recalled more often than associations from the third position (*t*_11_ = 6.27, *p* < 0.001).

In the alternating blocks of the behavioral experiment, participants recalled on average 69.47% (*SD* = 23.21%) of the correct word-scene associations in cued-recall (CR) blocks. They further recognized 90.27% (*SD* = 10.74%) of intact associations (Hits) and erroneously named 12.40% (*SD* = 14.38%) of rearranged associations intact (False Alarms) in an associative-recognition (AR) blocks. Performance in CR (i.e. percent correct responses) and in AR (i.e. percent Hits minus percent False Alarms) was compared with a 2×3 ANOVA. This revealed a significant main effect of condition (*F*_1, 23_ = 38.30, p < 0.001), driven by a better performance in the associative-recognition blocks (*t*_23_ = 6.189, *p* < 0.001) and a significant factor position (*F*_1.84, 42.24_ = 1.145, *p* = 0.002, interaction condition with position *n.s.*). This was driven by a slightly better performance in the cued-recall condition, for associations that were learned in the second position of a video-episode (ANOVA: *F*_1.58,36.24_ = 2.794, *p* = 0.086, position 1 vs. 2: *t*_23_ = −2.804, *p* = 0.02, position 2 vs. 3: *t*_23_ = 1.961, *p* = 0.062) and a worse performance in associative-recognition for associations that were learned in the third position (ANOVA: *F*_1.86,42.68_ = 5.552, *p* = 0.008, position 2 vs. 3: *t*_23_ = 3.879, *p* < 0.001, all other *ps* > 0.14).

In the MEG experiment subjects remembered on average 63.54% (*SD* = 11.768%) of associations, excluding guesses. After preprocessing on average 200.348 trials (*SD* = 38.645) remained for known correct associations and an additional 116 trials (*SD* = 39.425) were guessed or incorrect responses.

### Reaction time in the behavioral experiment (including correct guesses)

The analyses of reaction times were repeated including those trials in which participants indicated that they had guessed the response. The 2x3 ANOVA of RTs revealed a significant main effect of condition (*F*_1.00, 23.00_ = 66.254, *p* < 0.001, log-RT: *F*_1.00, 23.00_ = 98.52, *p* < 0.001) driven by overall faster reactions in the associative-recognition condition (*t*_23_ = −8.14, *p* < 0.001, log-RT: *t*_23_ = −9.619, *p* < 0.001). A significant main effect of scene-position (*F*_1.90, 43.67_ = 5.304, *p* = 0.010, log-RT: *F*_1.87, 43.09_ = 2.823, *p* = 0.074) and the interaction of scene-position with retrieval-condition (*F*_1.96, 45.11_ = 5.041, *p* = 0.011, log-RT: *F*_1.89, 43.39_ = 5.771, *p* = 0.007) were both due to a strong forward replay effect in the cued-recall condition (ANOVA: *F*_1.80, 41.36_ = 8.796, *p* = 0.001, log-RT: *F*_1.64, 37.82_ = 8.304, *p* = 0.002). Specifically, associations that were learned in the first scene-position of a video-episode (mean RT = 2.5 sec) were recalled on average 132ms faster than associations that were learned in the second scene-position (*t*_23_ = −1.752, *p* = 0.047, log-RT: *t*_23_ = −2.127, *p* = 0.022). Associations that were learned in the second scene-position (mean RT = 2.617 sec) were recalled on average 170ms faster than associations that were learned in the third scene-position (*t*_23_ = −2.864, *p* = 0.004, log-RT: *t*_23_ = −2.539, p = 0.009).

In the AR condition, subjects performed the exact same encoding task, which also required source-memory. Importantly, no differences in reaction times were evident between associations that were learned in the first, second or third position during encoding (ANOVA: *F*_1.44, 33.09_ = 0.185, *p* = 0.759, log-RT: *F*_1.52, 35.05_ = 0.591, *p* = 0.515, pairwise comparisons of positions: all *p*s > 0.5, Bayes-Factor supporting the null Hypothesis: position 1 vs. 2, *BF*_01_ = 3.771, position 2 vs. 3, *BF*_01_ = 4.466, position 1 vs. 3, *BF*_01_ = 4.504, log-RT: all *ps* > 0.39, position 1 vs. 2, *BF*_01_ = 3.688, position 2 vs. 3, *BF*_01_ = 3.317, position 1 vs. 3, *BF*_01_ = 4.048).

### Reaction times in the behavioral pilot

In the behavioral pilot experiment, participants associated word-cues with one of three scenes within video-episodes (Fig. 1a). Four continuous video-episodes each comprised of three individual scenes. A trial unique word-cue appeared in one scene during a video-episode. After a brief distractor task (Fig. 1b) a cued-recall task (Fig. 1d, top-left) was conducted where participants were presented with the word cues. Their task was to recall the scene-position that was associated with the word-cue as quickly as possible. After that, participants indicated which video-episode out of four was associated with the word.

Faster reaction times to associations that were associated with early position compared to later positions were observed (ANOVA: *F*_1.40, 15.41_ = 4.257, *p* = 0.045, ANOVA of log-transformed RTs: *F*_1.58, 17.38_ = 4.903, *p* = 0.027). On average, reaction times (RT) to first scene-positions were faster than RTs to second scene-positions (2.044 vs. 2.212 sec., *t*_11_ = −3.558, *p* = 0.005, log-RT: *t*_11_ = −3.626, *p* = 0.004), and trended to be faster than for third scene-positions (2.221 sec, *t*_11_ = −2.05, *p* = 0.065; log-RT: *t*_11_ = - 2.227, *p* = 0.048). RTs for second scene-positions were only numerically, but not significantly, faster compared to third scene-positions. These results suggest that memory replay is forward and compressed. During encoding, individual scenes of each video-episode lasted 2 seconds. During retrieval, however, subjects took on average 167.8ms longer to recall an association from the second scene-position and an additional 10ms longer to recall an association from the third scene-position. Importantly, these effects cannot be explained by material specific differences between positions, because balancing pilot experiments ensured that there were no differences in RTs when the scenes from position 1, 2 and 3 were associated with a word-cue in isolation (see Online Methods).

### Broad decreases in oscillatory power accompany successful memory reinstatement

Successful memory reinstatement was associated with strong and sustained decreases in oscillatory power. Successfully remembered associations were those trials, in which subjects knew that they had identified the correct scene and the correct video-episode. Those trials were contrasted with the trials in which subjects either indicated a guess, or in which they selected the wrong scene-position and/or video-episode. A broad cluster emerged, in which oscillatory power was significantly lower when memory-retrieval was successful (*p*_cluster_ < 0.001, Supplemental Fig. 1, middle). This cluster included a sustained power-decrease in the lower alpha band. In a series of post-hoc t-tests, the same contrast was now tested on averaged oscillatory power across time and sensors. Inspection of t-values confirmed a local peak at 8 Hz (*t*_22_ = −3.367, *p* = 0.001, Supplemental Fig. 1, right) which we previously linked to replay during episodic memory reinstatement ^8^. In order to derive the topography for the average power decrease at 8 Hz across time, a separate t-test was computed on every sensor. Maximal t-values were located over central sensors extending over right parietal sensors. The average power at 8Hz was next contrasted at every virtual sensor, resulting in an estimate of the spatial extent of power decreases in source space. Bilateral central and occipito-parietal areas as well as the medial temporal lobe displayed power decreases at this frequency (Supplemental Fig. 1, left). These findings replicate our previous findings of broad power decreases with a sustained decrease at 8Hz, in a paradigm that prompts subjects to replay a dynamic stimulus from memory.

**Supplemental Fig. 1:**
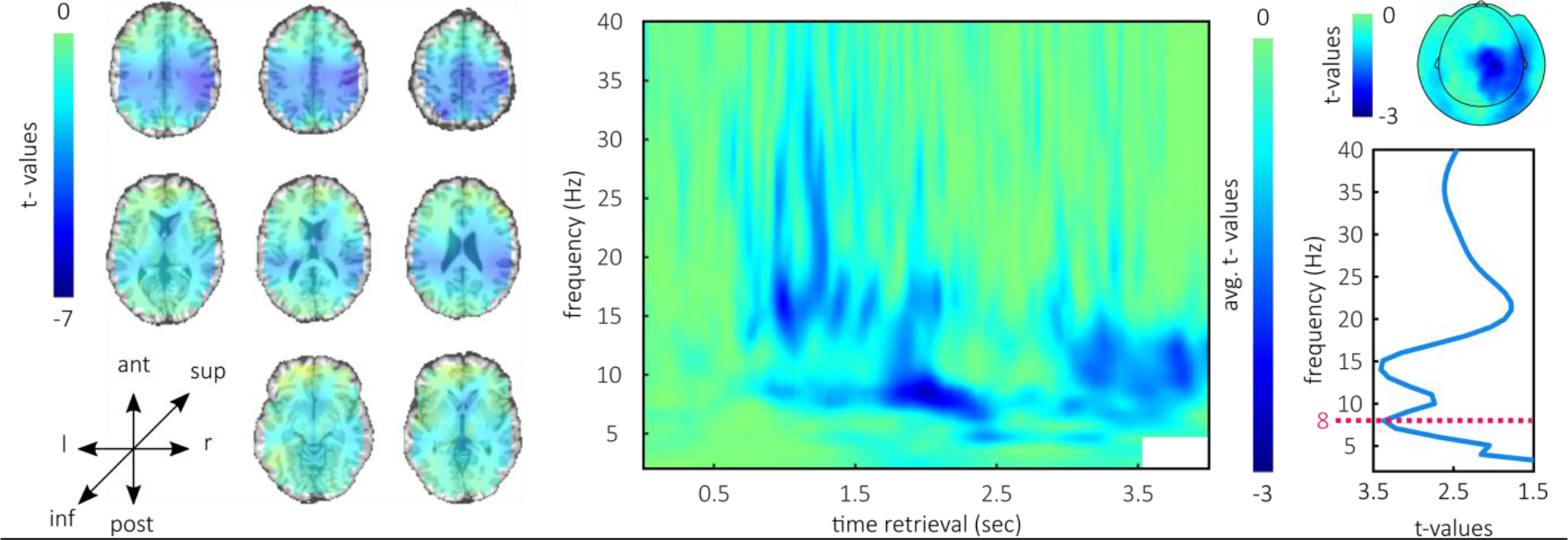
Oscillatory correlates of successful memory. Successful memory was associated with broad decreases in oscillatory power. The middle panel shows the t-values averaged across time and sensors within the significant cluster. Low frequencies displayed a sustained effect over time. T-tests of the average power decrease across time and sensors expressed two local peaks in t-values, at 8 and 14 Hz (right panel). The topography of t-tests for the 8 Hz frequency at every sensor included central sensors and extended over right parietal sensors (right panel, top). Source reconstructions of the average power at 8Hz revealed power decreases on bilateral central and occipito-parietal areas as well as the medial temporal lobe.

### Time course of replay and further evidence for forward replay

We further investigated the time-course of reinstatement of the video-episodes. To this end, we computed a t-test of content-specificity at every time point during retrieval; precisely we assessed the average content-specificity across the whole ROI (Supplemental Fig. 2a). Three peaks emerged at 442ms (*t*_22_ = 2.363, *p* = 0.014), 1042ms (*t*_22_ = 2.022, *p* = 0.028) and at 2163ms (*t*_22_ = 2.258, *p* = 0.017). Interestingly the last peak corresponds roughly to the period in which we observed average reaction times in the behavioral experiments.

Next, we further pursued forward replay. Following the behavioral results, we predicted that subjects replay overall more of the video-episodes when they have to recall later scene-positions (Supplemental Fig. 2b, left). We reasoned that subjects accumulate evidence in a forward direction, until the correct association is identified. In the behavioral experiments, this would cause the increase in RT for associations that were learned later during encoding. Hence, in analogy to the analysis of the behavioral experiment, we split the retrieval trials according to the remembered scene-position. We assessed the average similarity to all sub-scenes in the corresponding video-episodes from encoding and compared it between trials: We contrasted trials in which an association from the first, second or third positional-scene of a video-episode was remembered. In this overall similarity to the corresponding video-episode should be higher, when participants recalled associations from later scene-positions. Specifically, if subjects stop replaying consecutive scenes, once they found the correct position of an association, then overall similarity should increase linearly with the to-be-remembered scene-position (i.e. similarity 1 < 2 < 3). In a first ANOVA there was no significant difference in similarity depending on the remembered scene-position (*F*_1.83, 40.31_ = 2.384, *p* = 0.109, linear contrast: *F*_1, 22_ = 3.63, *p* = 0.07). There was, however, significantly less similarity to encoding, in trials in which subjects remembered an association from the first positional-scene compared to trials in which the third scene was recalled (*t*_22_ = −1.909, *p* = 0.035, Supplemental Fig. 2b). Note that this is equivalent to testing the hypothesis of a linear increase (i.e. directed linear contrast) with planned contrasts in the ANOVA.

**Supplemental Fig. 2:**
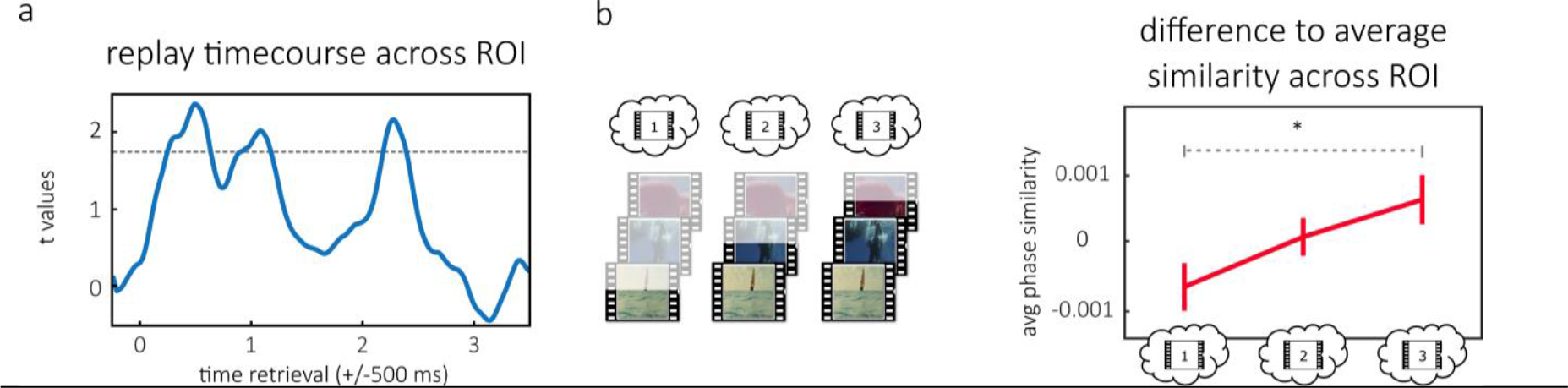
Time course of reinstatement and forward replay in ROI. **(a)** Time course of replay tested across the ROI. Three peaks are evident at 442ms, 1042ms and 2163ms. **(b)** Illustration of forward replay in which an association from the first, second or third scene was remembered (left) and difference to average similarity to encoding across the ROI (right, error bars are standard error of the mean). This shows higher similarity to encoding when associations from the third vs. first scene were remembered.

### Reinstatement of encoding patterns and further evidence for forward replay

Overall we found a cluster of significant evidence for the reactivation of phase patterns from encoding during retrieval for hit trials (Hits; *p_cluster_* = 0.034; Supplemental Fig 3a, Supplemental Fig 3b for unmasked maps of t-values). In this, we wanted to assess how much each sub-part from the video-episodes contributed to this effect. To this end, we computed a series of post-hoc t-tests for every encoding time-window. We obtained the highest t-values for the reinstatement of earlier time-windows during encoding (Supplemental Fig 3a, right).

If participants start to replay from the beginning of a video-episode and typically progress until they have the correct word-scene association in memory (compare Supplemental Fig 2b, left), then early time windows from encoding should be reactivated more often and more thoroughly (see above), therefore this pattern is consistent with forward replay.

**Supplemental Fig. 3:**
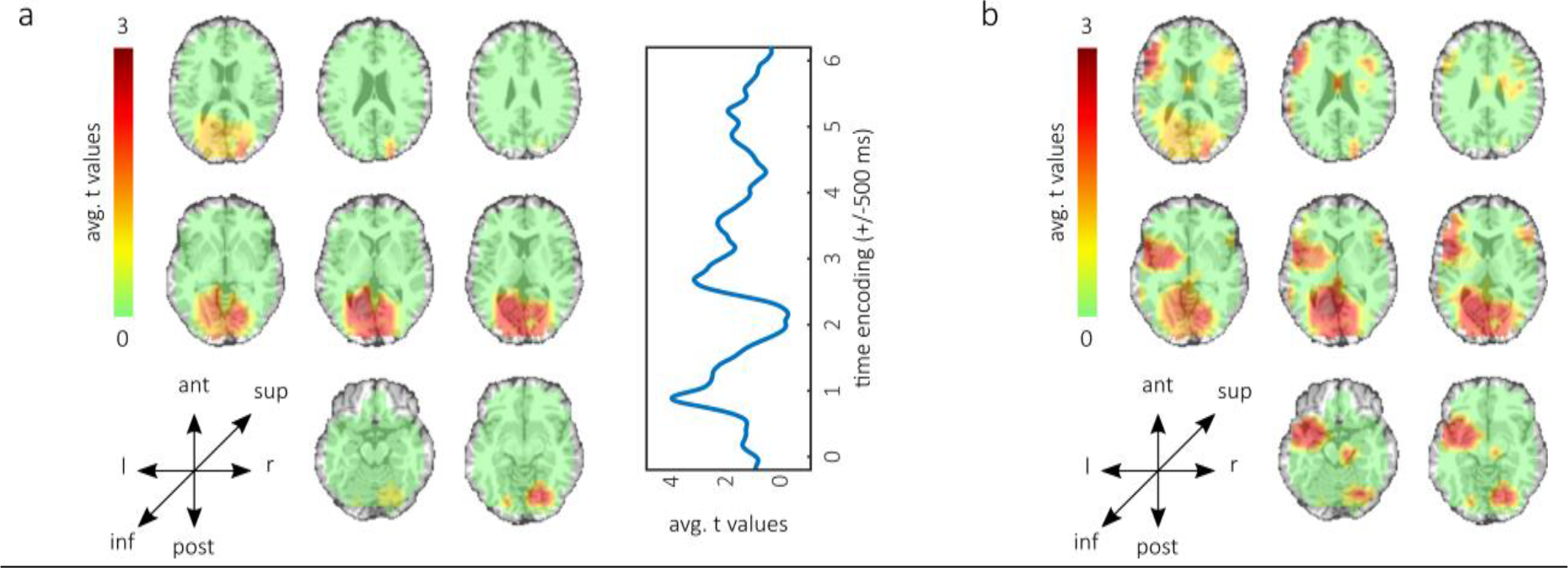
Content specific pattern reinstatement. **(a)** Cluster of significant reactivation of phase-patterns from encoding for successfully remembered associations (left) and contribution to effect (right). Early encoding windows express the highest t-values and contribute more to the effect than later ones. (**b)** Unmasked map of t-values for reactivation of phase-patterns from encoding in successfully remembered trials.

